# DeepFrag: A Deep Convolutional Neural Network for Fragment-based Lead Optimization

**DOI:** 10.1101/2021.01.07.425790

**Authors:** Harrison Green, David R. Koes, Jacob D. Durrant

## Abstract

Machine learning has been increasingly applied to the field of computer-aided drug discovery in recent years, leading to notable advances in binding-affinity prediction, virtual screening, and QSAR. Surprisingly, it is less often applied to lead optimization, the process of identifying chemical fragments that might be added to a known ligand to improve its binding affinity. We here describe a deep convolutional neural network that predicts appropriate fragments given the structure of a receptor/ligand complex. In an independent benchmark of known ligands with missing (deleted) fragments, our DeepFrag model selected the known (correct) fragment from a set over 6,500 about 58% of the time. Even when the known/correct fragment was not selected, the top fragment was often chemically similar and may well represent a valid substitution. We release our trained DeepFrag model and associated software under the terms of the Apache License, Version 2.0. A copy can be obtained free of charge from http://durrantlab.com/deepfragmodel.

## 2 Introduction

Drug discovery is benefiting from an upsurge in machine-learning approaches for tasks such as binding-affinity prediction [1–3], virtual screening [4–7], and QSAR [8]. Massive molecular datasets have enabled data-driven models that outperform handcrafted algorithms in nearly all applications. As these powerful new approaches come of age, they are increasingly used to augment the drug-discovery pipeline and reduce the time and cost of developing new pharmaceuticals.

While discriminative models for drug discovery have been studied extensively, the problem of molecular generation remains challenging. In the field of computer vision, generative models such as recurrent neural networks (RNNs) and generative adversarial networks (GANs) have had great success in performing tasks such as realistic image synthesis [9] and style transfer [10]. Naturally one wonders if this same technology can be applied to molecular synthesis. Several recent works in generative molecular modeling have demonstrated that it is possible to generate libraries of SMILES strings with desired properties [11] as well as 3D ligand pharmacophore-type maps from a given receptor pocket [12].

However, the field still faces some challenges. First, it is difficult to enforce the generation of valid molecular structures. Models that produce SMILES strings may contain grammatical errors, and 3D molecular shapes tend to be blurry and lacking in detail. Second, the inner workings of generative models are difficult to interpret. Failure cases are hard to diagnose both during training and inference. Finally, while it is clear how to evaluate a regressive model (L2 loss for example), it is not entirely clear how to quantitatively evaluate a generative model. The current best practice is to demonstrate “enrichment” for some metric (e.g., QED) compared to a random baseline [13].

In this paper, we address some of these limitations by restructuring the question of molecular generation as a type of classification problem. Specifically, we propose a new “fragment reconstruction” task where we take a ligand/receptor complex, remove a portion of the ligand, and ask the question “what molecular fragment should go here?” To successfully answer this question, a machine-learning model must consider the surrounding receptor pocket and the intact portion of the ligand. This task is immediately applicable to lead optimization, which seeks to improve the binding of a known ligand by swapping and/or adding molecular fragments. It also represents an important step towards fully *de novo* drug design.

We here demonstrate that a 3D convolutional network trained on experimentally derived crystal-structure data can select a missing fragment with >57% accuracy from a set of more than 6,500 fragments. Even when the network does not predict the “correct” answer, the top predictions are often chemically similar and may well represent plausible substitutions. We release our trained model and associated software under the terms of the Apache License, Version 2.0. A copy can be obtained free of charge from http://durrantlab.com/deepfragmodel, where interested users can also find a link to a Google Colaboratory Notebook [14] for testing.

## 3 Materials and Methods

### 3.1 Training Datasets: Receptor/Ligand Complexes

Our goal was to train a supervised model to complete a 3D structure of a receptor-bound partial ligand (“parent”) with a molecular fragment, such that the resulting composite molecule (parent + fragment) is highly complementary to the receptor. To this end, we first constructed a library of (*receptor/parent*, *fragment*) tuples, where each fragment is a well-suited choice for the corresponding receptor/parent complex. We assembled this library from the Binding MOAD dataset [15], which includes experimentally derived structure data for 38,702 receptor/ligand complexes (Figure 1A).

**Figure 1:**
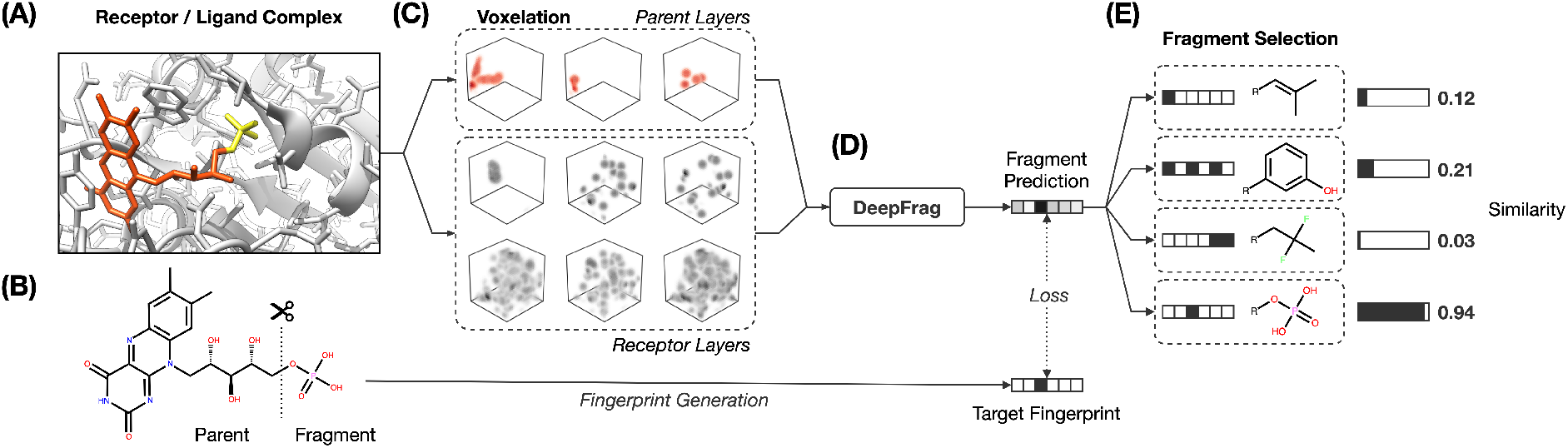
DeepFrag workflow. (A) Receptor/ligand complex. Example “parent” and “fragment” portions of the ligand are highlighted in orange and yellow, respectively. (B) The ligand is cut along a single bond to separate the parent and fragment. (C) Atoms from the receptor and parent are converted to 3D voxel grids (density channels). (D) Density channels are concatenated and fed to the DeepFrag model, which predicts a fragment fingerprint vector. (E) The fingerprint vector is compared against a fingerprint library (label set) to generate predictions.

#### 3.1.1 Data Pre-processing

The crystal structures contained water molecules as well as crystallographic additives (e.g., buffers, salts). The Binding MOAD specifically catalogues which ligands are biologically relevant. We used these annotations to strip irrelevant artifacts before generating fragments. Additionally, some complexes contain several bound ligands per receptor. In these cases, we isolated each ligand as a separate receptor/ligand complex.

#### 3.1.2 Ligand Fragmentation

For each receptor/ligand complex, we generated multiple (*receptor/parent*, *fragment*) training examples by iterating over all ligand single bonds and performing a cut (Figure 1B). We retained the resulting example if it satisfied the following criteria:

- the cut split the ligand into two disconnected pieces (e.g., cutting a ring was not permitted)
- the smaller piece contained at least one heavy atom
- the smaller piece had a molecular weight < 150 Da
- the connection point between the parent and fragment was within 4 Å of a receptor atom

When these conditions were met, we labeled the smaller and larger pieces of the ligand the *fragment* and *parent*, respectively. All fragments were standardized using MolVS transformation rules [16]. Specifically, for each fragment, we neutralized the charge and generated a canonical tautomer. By constraining our dataset to only fragments located near the receptor surface, we ensured that they were likely to form interaction(s) with their respective receptors. There were ultimately 308,689 (*receptor/parent*, *fragment*) tuples that met these criteria.

#### 3.1.3 Data Splits

The examples were partitioned into three sets (*TRAIN*, *VAL*, and *TEST*), in the approximate ratio 60/20/20. We trained models on the *TRAIN* set and used the *VAL* set to monitor performance, tune hyperparameters, and prevent over-fitting. We report final results on the withheld *TEST* set.

Some protein targets are repeated in the Binding MOAD. To ensure our model generalized to unseen receptors, we created a three-way “vertical split” so that homologous targets were not shared between the *TRAIN*, *VAL*, or *TEST* sets. The Binding MOAD provides 90% similarity families that we used to determine if two targets were homologous.

Similarly, many ligands bind to multiple targets, and some metabolites such as ATP occur very frequently. To prevent the model from simply memorizing common ligands, we further ensured that ligands (in addition to receptors) were not shared between the *TRAIN*, *VAL*, and *TEST* sets. We examined the previous “vertical split” dataset and identified ligands shared across multiple splits by comparing canonical SMILES strings. Each shared ligand was then randomly assigned to one of the sets, and examples from the other set(s) were discarded.

### 3.2 Receptor/Parent Complexes as Voxel Grids

We converted the structures of each receptor/parent example into a 3D voxel grid (Figure 1C). To generate each grid, we placed a virtual box in the receptor pocket, centered on the fragment/parent connection point (i.e., the location of the parent atom from which the fragment should branch). This section describes the specific parameters we considered when generating these grids.

#### 3.2.1 Grid Widths and Resolutions

We used cubic grids. The grid width, *w*, is defined as the number of grid points in each dimension, such that there are *w*^3^ points total. The spacing in Ångstroms between the grid points, *s*, is called the grid resolution. The physical length of the grid is thus *ws* along each dimension, and the volume is *w*^3^*s*^3^.

#### 3.2.2 Voxelation methods

Our grid representation requires that each atom contribute “density” to neighboring grid points that fall within a certain distance (the atom-influence radius, *r*). We considered several functions for calculating these densities (Figure 2):

1. SPHERE: Set all grid points within *r* to 1 (spherical boolean).
2. CUBE: Set all grid points within *r* along the *x*, *y*, or *z* axis to 1 (cubical boolean).
3. POINT: Set only the nearest grid point to 1 (as in ref. [2]).
4. SMOOTH: Set nearby grid points per a continuous, piecewise function (as in ref. [5]), defined as:

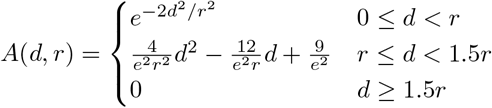

where *d* is the distance from the grid point to the atom center. Note that in the present work, *r* is the same for all atom types and so is not equivalent to an atomic radius.
5. SMOOTH-2: Set nearby grid points per the exponential part of SMOOTH:

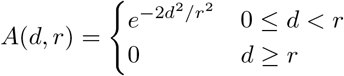
6. LJ: Set grid points per the repulsive component of a Lennard-Jones potential (as in ref. [17]), defined as:

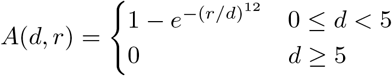

**Figure 2:**
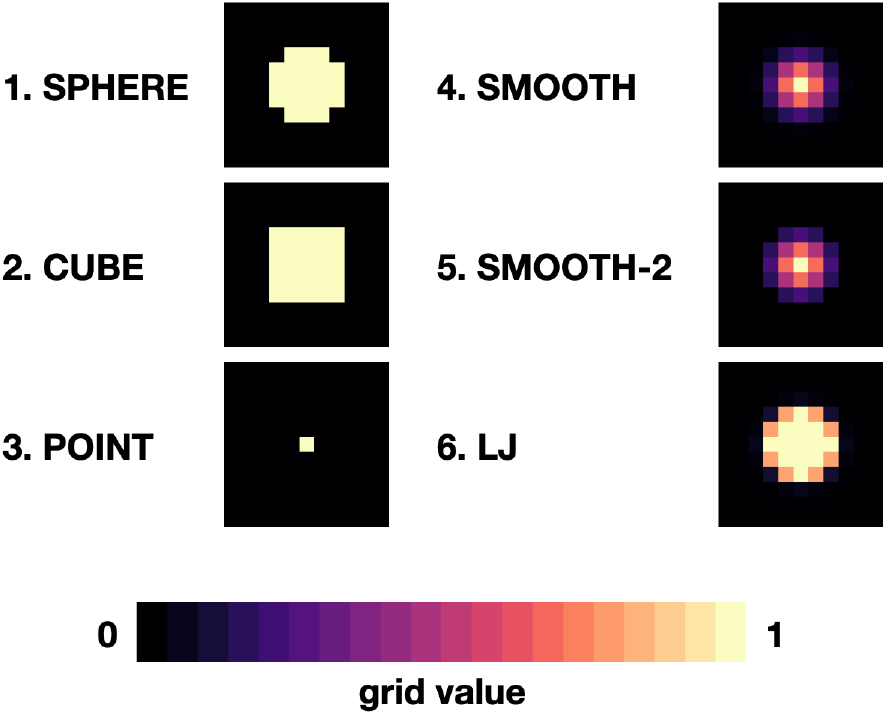
Illustration of different grid-voxelation methods in 2D.

#### 3.2.3 Grid Layers

To distinguish different atom types, we considered several “atom-channel” (i.e., grid-layer) definitions. In all cases, atoms from the receptor and parent always contributed to different grid layers, allowing our model to distinguish between the two.

In the *flat* definition, we assigned all atoms to a single channel. This method serves as a baseline comparison and is analogous to converting an image to grayscale before training an image classifier. The *flat-h* definition includes hydrogen atoms, but *flat* does not.

In the *simple* definition, we assigned atoms to separate layers based on atomic number. We assigned the most common atoms to individual layers and aggregated all other atoms in a separate “other” layer. For ligands, the most common atoms are carbon, nitrogen, and oxygen. For receptors, we also include a separate sulfur layer. The *simple-h* definition further includes a hydrogen layer.

We also experimented with high-level descriptors for receptor atoms. In the *meta* definition, we assigned each atom to one or more channels based on the following properties: aromatic, hydrogen-bond donor, hydrogen-bond acceptor, partial positive charge, partial negative charge, and occupancy (any atom). For example, an aromatic carbon atom with a partial positive charge would be present in the aromatic, partial positive, and occupancy layers. In the *meta-mix* description we combine the *meta* layers and the *simple-h* layers.

#### 3.2.4 Data-Handling Optimizations

We used two optimizations to accelerate data handling. First, to load the receptor/parent data faster, we stripped all irrelevant information from the source PDB files and saved only atomic coordinates and atom types to a packed HDF5 file. During training, we loaded this entire dataset into memory for rapid access, thereby drastically decreasing training startup time.

Second, to convert this data to a voxel grid, we developed a GPU-accelerated grid-generation routine using Numba, a high-performance Python just-in-time compiler [18]. As others have noted [19], a GPU-accelerated approach reduces grid-generation wall-time substantially (more than 10x in our case). We therefore opted to generate all grids on the fly each time they were needed (i.e., once per epoch), without ever saving them to the disk or storing them long-term in memory. As part of the grid-generation process, we randomly rotated each example. The model was thus trained on differently rotated grids every epoch, a form of data augmentation aimed at encouraging rotation-invariant learning.

### 3.3 Representing Fragments as Vectors

We converted the fragments of the *TRAIN*, *VAL*, and *TEST* sets to vectors by applying the RDKFingerprint algorithm [20] provided by the RDKit library [21]. RDKFingerprint is an implementation of a Daylight-like topological fingerprint, which enumerates subpaths in a molecule and computes hashes. Each hash seeds a pseudo-random number generator, which is used to randomly set bits in the fingerprint bitstring. Molecules with matching subpaths share common bits. Specifically, we allowed the subpath enumeration to consider all paths of size ≤ 10 to differentiate between larger fragments (e.g., alkanes). We folded the final bitstring to a size of 2048. The generated fingerprints contained only 0’s or 1’s, but we allowed our model to predict continuous vectors with each element in the range [0, 1] (enforced with a sigmoid activation layer).

### 3.4 DeepFrag Model

#### 3.4.1 Architecture

We used a deep 3D convolutional neural network to predict appropriate RDKFingerprint fragment vectors from voxel-grid representations of receptor/parent complexes (Figure 1D). The model consists of several repeated blocks of 3D convolution layers followed by a global average pooling layer and several fully connected layers. Each convolution block starts with a batch normalization layer and ends with a 3D max pooling layer (except the last block). In the fully connected section, we use dropout layers to prevent overfitting. All intermediate activations are ReLU except for the last layer, which uses a sigmoid activation to map values into the range (0,1). The final “optimized” model architecture is shown in Figure 3.

**Figure 3:**
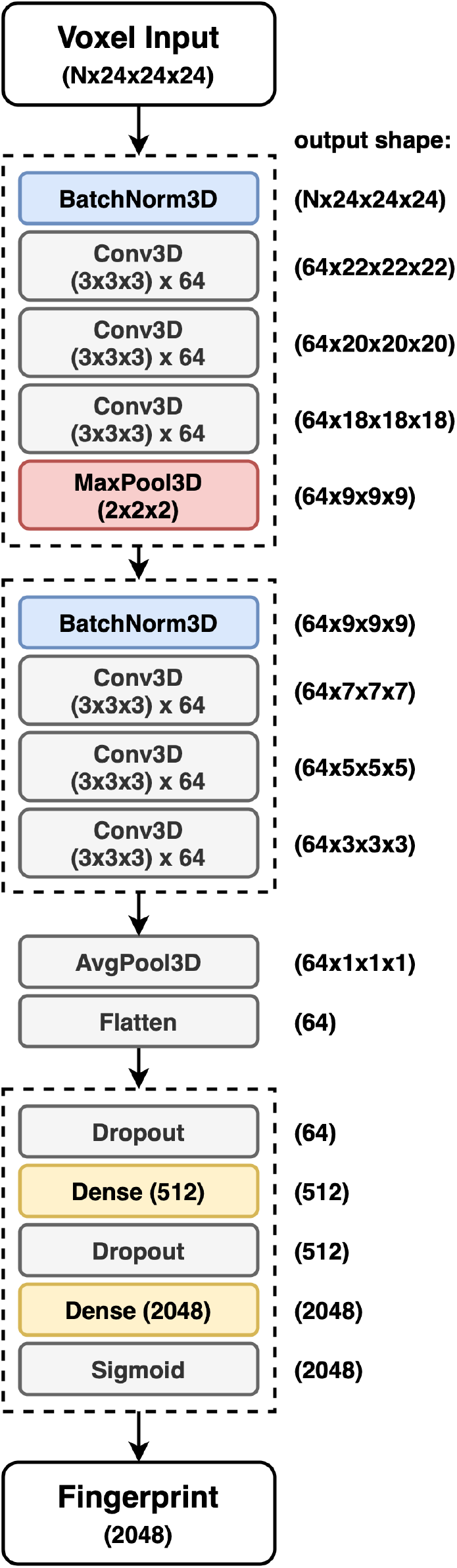
Final DeepFrag convolutional neural network architecture. The input tensor consists of concatenated atom-wise channels (grids) from the parent and receptor (*N* = total number of atom channels). The output tensor contains a raw fragment-fingerprint prediction. For this model, blocks = [64,64] and fc = [512] (see Supporting Information).

#### 3.4.2 Training Details

Models were implemented in PyTorch [22]. We used the Adam optimizer [23] with the default momentum schedule. For each epoch, we randomly sampled batches from the *TRAIN* set and computed predicted fingerprint vectors after applying random grid rotations. The loss was computed as the average cosine distance between the predicted fragment vectors and the vectors of the corresponding *correct* (known) fragments for each batch (Equation 1).

To monitor training, we randomly sampled a subset of the *VAL* set after each epoch and computed an average validation loss using the same loss function. We only saved new model checkpoints when the validation loss reached a new global minimum. Training continued until we observed convergence. To accelerate training, we trained models using either *NVIDIA Titan X* or *NVIDIA GTX1080* GPUs.

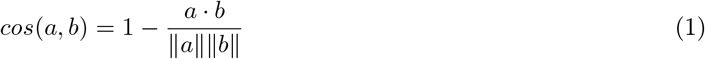

Equation 1: Cosine distance between vectors *a* and *b*. A distance of 0 represents parallel vectors. A distance of 1 represents orthogonal vectors.

### 3.5 Label Sets, Fragment Selection, and TOP-k Accuracy

The model itself predicts a vector in RDKFingerprint space, not a human-interpretable fragment representation. To map an output vector to specific fragments, we compute the cosine distance (Equation 1) between the predicted vector and each vector in a large library of RDKFingerprints corresponding to known fragments (Figure 1E), which we call a “label set.” Using these distances we can rank fragments by similarity to the predicted vector and identify the “closest match.” The label set is not necessarily the same set of fingerprint vectors used to train, validate, or test the model. Rather, it is the set of all fragments from which the model can ultimately choose.

If the label set contains fragments known to be *correct* (i.e., if it includes the fragments of the *VAL* and/or *TEST* sets), it is possible to quantitatively evaluate a model’s accuracy. We consider TOP-k accuracy, for *k* ∈ {1, 8, 64}. For example, TOP-8 accuracy represents the probability of finding the *correct* fragment in the set of the top eight predicted fragments.

### 3.6 Hyperparameter Tuning

We optimized hyperparemeters in several phases. Early-stage efforts focused on identifying suitable model and grid-density parameters (see Supporting Information). We here describe our subsequent efforts to identify suitable voxelation and grid-layer parameters. For these steps, we trained on examples in the *TRAIN* set and evaluated TOP-k accuracy on examples in the *VAL* set. We merged the fragment vectors from the *TRAIN* and *VAL* sets into a combined label set, which we used for fragment selection.

We trained each model variant for 15 epochs (approximately 20 hours) on a GPU, achieving accuracy roughly 80-90% of the eventual maximum. During early experiments, we found that this shortened training cycle was sufficient to assess relative validation accuracy between model variants. That is, in general if a model performed better after 15 epochs, it also performed better after full convergence.

Using suitable model and grid-resolution parameters identified previously (see Supporting Information), we explored the effect of varying the grid-layer definitions of the parent and receptor atoms. When the parent-atom definition was varied, the receptor-atom definition was fixed as *simple*, and vice versa. We trained each of the 9 model variants in triplicate and report the average validation accuracy after 15 epochs.

Additionally, we experimented with changing the shape and size of atom densities by performing a comprehensive grid search. In this experiment, we fixed the parent and receptor typing schemes as *simple* and varied the voxelation method and atom-influence radius (Table 2).

**Table 1:** Grid-layer (atom typing) descriptions. Note that the *simple* and *simple-h* variants for the receptor include a separate sulfur layer. Otherwise, the parent and receptor typing schemes are identical. Additionally, the *meta* and *meta-mix* descriptions are used only for the receptor.

| Parent Typing | Layers |
| --- | --- |
| flat | single channel, no hydrogen |
| flat-h | single channel, including hydrogen |
| simple | [C, N, O, (other)] |
| simple-h | [H, C, N, O, (other)] |
| Receptor Typing | Layers |
| flat | single channel, no hydrogen |
| flat-h | single channel, including hydrogen |
| simple | [C, N, O, S, (other)] |
| simple-h | [H, C, N, O, S, (other)] |
| meta | [aromatic, H-don, H-acc, pos, neg, occupancy] |
| meta-mix | *meta* + *simple-h* |

**Table 2:**
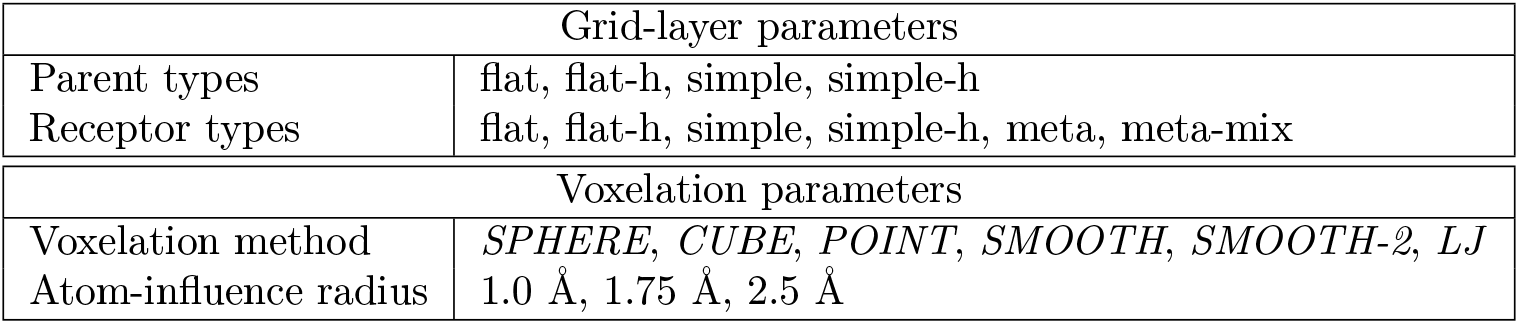
Select hyperparameters varied during our tuning protocol. During the grid-layer experiment, the receptor typing scheme was fixed at *simple* while the parent typing scheme was varied, and vice-versa.

### 3.7 Final Model: Training to Convergence

Once we settled on a set of effective hyperparameters, we trained an “optimal” model, which we call DeepFrag. We trained the final model to convergence (5 days on a GPU) and evaluated its TOP-k accuracy on the withheld *TEST* set, using as a label set the fragment vectors of the *TRAIN*, *VAL*, and *TEST* sets. Training to convergence required substantial computer resources. But using DeepFrag to prospectively evaluate a single receptor/parent complex (i.e., at inference time) requires only a few grid-generation steps that can easily run on a CPU.

Using the fully converged model, we also explored the effect of test-time augmentation on model accuracy. We evaluated the effect of sampling multiple random rotations per example and averaging the predicted fingerprints to generate a multi-rotation, ensemble prediction.

## 4 Results and Discussion

The fragment-reconstruction task we here introduce aims to complete a partial, receptor-bound ligand (“parent”) by adding a new molecular fragment such that the combined molecule is highly complementary to the given receptor. For the purpose of this work, a fragment is a terminal ligand substructure with a mass less than 150 Da. The cutoff choice of 150 Da is relatively arbitrary; this value allows for a large variety of fragment types without including overly complex structures. The fragment may be an explicit functional group with known behavior such as a hydroxyl or phenyl group, but this is not a requirement.

The intuition behind this task is that fragment selection is a function of the local receptor environment and the existing ligand scaffold (parent). Therefore, it should be possible to learn a model *f*(*receptor, parent*) = *fragment*. We here introduce just such a model and show that it can be used to implicitly rank a set of candidate complementary fragments. We expect models such as these to be useful for lead optimization (e.g., to generate congeneric series of small-molecule ligands with improved binding affinities). The same model could be used indirectly as a way to evaluate the importance of each group in an existing ligand by removing and re-predicting existing fragments.

### 4.1 Generating and Representing Molecular Data

#### 4.1.1 Assembling a Dataset of (*Receptor/Parent*, *Fragment*) Tuples

Ideally, we would like to train a model to predict the single, optimal fragment for any receptor/parent pair. But given that there are roughly 10^60^ drug-like molecules [24], identifying optimal fragments for training is impracticable. We instead trained on datasets derived from the Binding MOAD [15] database (Table 3), which currently includes experimentally derived structural data for 38,702 receptor/ligand complexes.

**Table 3:** Fragment dataset details. *Receptors*: number of unique receptor targets per split. *Ligands*: number of unique ligands per split (determined by SMILES comparison). *Unique Fragments*: number of unique molecular fragments per split. *Examples*: total number of (*Receptor* /*Parent*, *Fragment*) tuples.

|  | TRAIN | VAL | TEST | ALL |
| --- | --- | --- | --- | --- |
| Receptors | 22,968 | 7,641 | 8,093 | 38,702 |
| Ligands | 10,284 | 3,677 | 4,101 | 18,062 |
| Unique Fragments | 4,654 | 2,070 | 2,328 | 6,522 |
| Examples | 185,198 | 55,221 | 68,270 | 308,689 |

We make the assumption that because each ligand in the dataset is known to bind the corresponding target, its structure must be to some extent optimized relative to a random molecule. It follows that each ligand fragment (i.e., substructure) is also at least somewhat optimized—especially those fragments that interact directly with the receptor. We therefore trained a model to reconstruct *correct* fragments from known active ligands, as a surrogate for training on *optimal* fragments.

#### 4.1.2 Voxelizing Receptor/Parent Complexes

We represented receptor/parent complexes as 3D voxel grids (tensors) similar to those described elsewhere [4, 5]. We chose a grid representation because the 3D local context is certainly critical for fragment binding. Converting molecular structures to voxel grids also allowed us to easily translate machine-learning techniques from other fields (e.g., computer vision). To learn a rotation-invariant model and prevent overfitting, each grid was randomly rotated each time it was used (i.e., once per training epoch).

#### 4.1.3 Converting Fragments to Fingerprint Vectors

We converted molecular fragments to vectors using the RDKFingerprint algorithm [20, 21]. This fingerprint can be generally seen as a vectorized description of the fragment’s topology, or structure. Our approach thus differs from a typical categorical classification task, where a model predicts a vector containing class scores or normalized class probabilities. In this typical formalization, classes must be fixed at training time (i.e., the model has no capacity to predict unseen classes), and any prior knowledge about class relationships is effectively stripped, precluding the training of a more generalizeable model. In our fragment prediction task, we expect structurally similar fragments to have similar binding properties; by training on fingerprint vectors, we enforce this prior knowledge.

### 4.2 Hyperparameter Tuning

We systematically explored a number of hyperparameter combinations to identify one well-suited for fragment prediction (see Supporting Information and Section 3.6). We found that the choice of hyperparameters can substantially impact both the speed of grid generation and the quality of information ultimately provided to the machine-learning model.

#### 4.2.1 Model Learning and Architecture

We explored the impact of varying parameters related to network architecture and training (Table S1). Our base model (Section 3.4.1) was inspired by related works that use 3D atomic densities with convolutional neural networks [5, 17]. We ultimately selected a learning rate of 0.0001, a batch size of 16, and a model architecture with two convolutional blocks of size 64 and a single fully connected layer of size 512 (Figure 3).

#### 4.2.2 Grid-Density Parameters

We also varied the grid width and resolution parameters used to generate the voxel grids. Both these variables impact the speed of grid generation and the amount of information provided to the network. The grid resolution (i.e., the distance between adjacent grid points) is particularly impactful. Low-resolution grids can be generated quickly, but high-resolution grids provide more information. Based on our hyperparameter search, we selected a grid width of 24 and a grid resolution of 0.75 Å (Table S1).

#### 4.2.3 Grid-Layer Parameters

In the same way a 2D image has three color channels (e.g., red, green, blue), our 3D grids have *N* atom channels (e.g., carbon, nitrogen, oxygen, etc.). It is tempting to construct a large array of features (e.g., to create separate layers for aliphatic and aromatic carbon atoms) in order to maximize information. But using too many layers can slow training and make the model harder to deploy because each feature must be separately computed during inference. On the other hand, too few layers may result in information loss and worse performance.

We tested several grid-layer schemes (Table 4) and ultimately settled on a straightforward approach based solely on the atomic elements of receptor (N, O, C, S, other) and parent (N, O, C, other) heavy atoms, without regard for atomic hybridizations (i.e., the *simple* definition for both the receptor and parent, see Section 3.2.3). Other works [5] have found that element-type channels are sufficient to achieve high performance on the related binding-affinity prediction task. Similarly, including hydrogen atoms appears to have little impact on affinity-prediction performance [25]. In our preliminary tests, we similarly saw little benefit to including hydrogen atoms (Table 4).

**Table 4:** Effect of varying grid layer descriptions on TOP-1 accuracy. For each parent grid-layer variant, the receptor grid-layer definition was fixed at *simple*, and vice-versa. Each model variant was trained in triplicate on the *TRAIN* dataset for 15 epochs. We report TOP-1 accuracy (%) and standard error on the *VAL* set.

| Parent Typing | Receptor Typing | TOP-1 Accuracy |
| --- | --- | --- |
| flat | simple | 45.87 ($\pm 0.44$) |
| flat-h | simple | 46.00 ($\pm 0.55$) |
| simple | simple | 50.57 ($\pm 0.25$) |
| simple-h | simple | 50.45 ($\pm 0.06$) |
| Parent Typing | Receptor Typing | TOP-1 Accuracy |
| simple | flat | 46.20 ($\pm 0.51$) |
| simple | flat-h | 47.27 ($\pm 0.83$) |
| simple | simple | 50.57 ($\pm 0.25$) |
| simple | simple-h | 50.10 ($\pm 0.58$) |
| simple | meta | 49.22 ($\pm 0.58$) |
| simple | meta-mix | 50.10 ($\pm 0.87$) |

Our straightforward grid-layer definition is particularly advantageous in terms of end-user usability. Most crystal structures lack the resolution required to correctly position hydrogen atoms, and computational methods for predicting protonation/tautomerization states are error prone. Similarly, predicting ligand-atom hybridizations from 3D coordinates is challenging. Because DeepFrag does not require this information, end users deploying DeepFrag in production do not need to guess at these atomic properties prior to use.

#### 4.2.4 Voxelation Parameters

We also varied the parameters used to convert atomic coordinates to voxel-grid densities (Table 5). The voxelation method controls the *shape* of atomic densities 2 and the atom-influence radius controls the *size* of atomic densities (Tables 2 and S1).

**Table 5:** Effect of voxelation method on TOP-1 accuracy (%). Each model variant was trained three times on the *TRAIN* dataset for 15 epochs. We report TOP-1 accuracy (%) and standard error on the *VAL* set.

|  | 1.0 Å | 1.75 Å | 2.5 Å | Overall |
| --- | --- | --- | --- | --- |
| 1. SPHERE | 49.12 ($\pm 0.48$) | 49.65 ($\pm 0.38$) | 48.50 ($\pm 0.43$) | 49.09 ($\pm 0.30$) |
| 2. CUBE | 49.27 ($\pm 0.55$) | 42.12 ($\pm 0.13$) | 39.18 ($\pm 0.13$) | 44.06 ($\pm 1.41$) |
| 3. POINT | 44.13 ($\pm 0.18$) | 43.27 ($\pm 0.48$) | 43.40 ($\pm 0.56$) | 43.64 ($\pm 0.28$) |
| 4. SMOOTH | 50.38 ($\pm 0.08$) | 49.00 ($\pm 0.06$) | 46.93 ($\pm 0.70$) | 48.74 ($\pm 0.56$) |
| 5. SMOOTH-2 | 51.13 ($\pm 0.61$) | 49.40 ($\pm 0.61$) | 46.23 ($\pm 0.74$) | 48.92 ($\pm 0.78$) |
| 6. LJ | 50.03 ($\pm 0.42$) | 48.68 ($\pm 0.88$) | 47.32 ($\pm 0.30$) | 48.68 ($\pm 0.50$) |
| Overall | 49.01 ($\pm 0.58$) | 47.13 ($\pm 0.76$) | 45.62 ($\pm 0.71$) | |

Various grid-voxelation methods can convert atomic coordinates and atom types into 3D density tensors. Simple boolean voxelation is fast to compute but may oversimplify the input representation, while more complex functions can retain high-resolution distance information at the cost of slower grid generation. Surprisingly, the specific method had little effect on overall prediction accuracy (Table 5), with the exception of the *CUBE* and *POINT* methods, which apparently fail to provide sufficient information to the model. We hypothesize that the *CUBE* method does not preserve atomic symmetries (especially as the atom-influence radius increases), and the *POINT* method is too sparse. As others have noted [5], the hard boolean cutoff in *SPHERE* and the smooth Gaussian in *SMOOTH* are remarkably competitive, even though the former does not preserve atomic-distance information.

We ultimately settled on the *SMOOTH-2* voxelation method (1.75 Å radius) because it is the fastest to compute of the distance-preserving spherical methods. However, our hyperparameter search did not reveal a clearly optimal voxelation method and atom-influence radius; indeed, different methods/radii produce similar results.

### 4.3 Training a Final Production Model

#### 4.3.1 Training to Full Convergence

After exploring different combinations of hyperparameters, we trained an “optimal” model to full convergence, using the examples of the *TRAIN* set. Training ran for approximately 50 epochs (5 days on a TITAN-X GPU). We call this fully converged, final model “DeepFrag.” For this production model, we used a learning rate of 0.0001, a batch size of 16, and the model architecture shown in Figure 3. We generated cubic grids of resolution 0.75 Å and width 24 points. Grid density was calculated using the *SMOOTH-2* voxelation method (atom-influence radius of 1.75 Å). We used the *simple* grid-layer definition (based solely on atom elements) for both the receptor and parent.

#### 4.3.2 Averaging the Fragment Vectors of Multiple Rotated Grids

Although all grids were randomly rotated once per epoch to encourage rotation-invariant training, we found that the model was still not perfectly robust to rotation. That is, otherwise identical grids with different rotations generated slightly different prediction vectors, and some rotations even generated poor predictions. We found that averaging the fingerprint predictions of multiple randomly rotated grids improved accuracy by “smoothing out” any rotation dependence of the converged model. In Table 6 we report the TOP-k test accuracy on the *VAL* set obtained by averaging *N*-rotations per sample for different values of *N*.

**Table 6:** Effect of multiple-rotation sampling on the accuracy of a fully converged model (roughly 50 epochs of training). Each entry represents the TOP-k *VAL*-set accuracy obtained by averaging *N*-rotations per sample.

| N | TOP-1 | TOP-8 | TOP-64 |
| --- | --- | --- | --- |
| 1 | 56.41 | 64.79 | 70.78 |
| 2 | 57.17 | 65.47 | 71.36 |
| 4 | 57.57 | 65.87 | 71.78 |
| 8 | 57.72 | 66.03 | 71.98 |
| 16 | 57.83 | 66.06 | 72.05 |
| 32 | 57.91 | 66.17 | 72.13 |

#### 4.3.3 Final Evaluation on the *TEST* Set

As a final evaluation of the fully converged DeepFrag model, we considered the examples of the *TEST* set. For each *TEST*-set tuple, we randomly rotated the associated receptor/parent voxel grid 32 times and used DeepFrag to predict 32 fragment vectors. In all cases, we averaged the 32 vectors to produce one vector per receptor/parent pair. To calculate TOP-k accuracies, we compared these averaged vectors to the set of all fragment vectors in the *TRAIN*, *VAL*, and *TEST* sets (the “*LBL-ALL*” label set). The *LBL-ALL* vector closest to the predicted vector was the *correct* vector 57.77% of the time (Table 7).

**Table 7:**
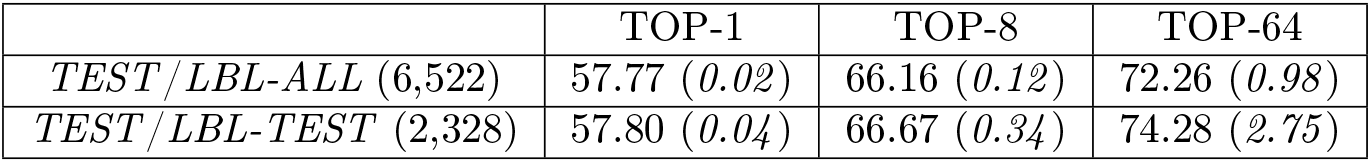
The TOP-k % accuracy, for *k* 1, 8, 64, of the final model (DeepFrag) evaluated on the withheld *TEST* set (68,270 examples), using the *LBL-ALL* label set (default). For reference, we also report the accuracy using the *LBL-TEST* set. The expected accuracy of an equivalent random model is given in parenthesis (italics).

### 4.4 Best Practices for Fragment Selection: Label Set Size

A large label set such as the *LBL-ALL* set (6,522 vectors) is advantageous in that it gives the model the freedom to select from a wider range of fragments. On the other hand, smaller sets may also have their advantages. For example, allowing DeepFrag to choose from a smaller label set could conceivably improve accuracy. Smaller sets comprised of easily synthesizeable fragments may also be critical when chemical synthesizeability is a concern. Even a few dozen fragments can cover a wide range of potential biochemical interactions.

To assess the impact of label-set size, we applied DeepFrag to all the (*receptor/parent*, *fragment*) tuples in the *TEST* set (68,270 tuples) and recalculated TOP-k accuracy using a much smaller label set comprised of only the fragment vectors present in our *TEST* set (*LBL-TEST*). Remarkably, the correct label was the closest roughly 58% of the time (Table 7), regardless of the label set used. Increasing the size of the label space by nearly threefold (*LBL-TEST* -> *LBL-ALL*) thus bears little cost on accuracy. In other words, model predictions are fairly precise; despite “cluttering” the label space with many more incorrect fragments, top predictions from the smaller space are generally also top predictions in the larger space.

### 4.5 Examples Demonstrating Effectiveness

#### 4.5.1 DeepFrag Generalizability

We were encouraged to see evidence that our model can generalize beyond the *correct* fragments used for training. The true *optimal* fragment for a given protein/parent pair is unlikely to ever be in any chosen label set. Generalizability ensures that the model can nevertheless predict chemically similar fragments that are well suited for a given binding-pocket region. Synthesizing and testing congeneric series of distinct compounds that each incorporate a different top-scoring fragment may enable the rapid discovery of optimized ligands with improved binding affinities.

To illustrate, we randomly selected six *TEST*-set (*protein/parent*, *fragment*) tuples and identified the eight *LBL-ALL* fragment fingerprints that were most similar to each predicted vector (Figure 4). The *correct* fragment was selected first in two of the six cases, and among the top five in another two cases. But it is telling that the other top-ranked fragments are often very plausible substitutes. Interestingly, the single-atom halogen fragments (*Cl, *Br, and *F) share no common fingerprint bits, yet the model learned to group them together. This result suggests that our model may be more predictive than even the TOP-k metric would suggest. It is entirely possible that in some cases where DeepFrag does not select the *correct* fragment, the selected fragment may in fact be superior.

**Figure 4:**
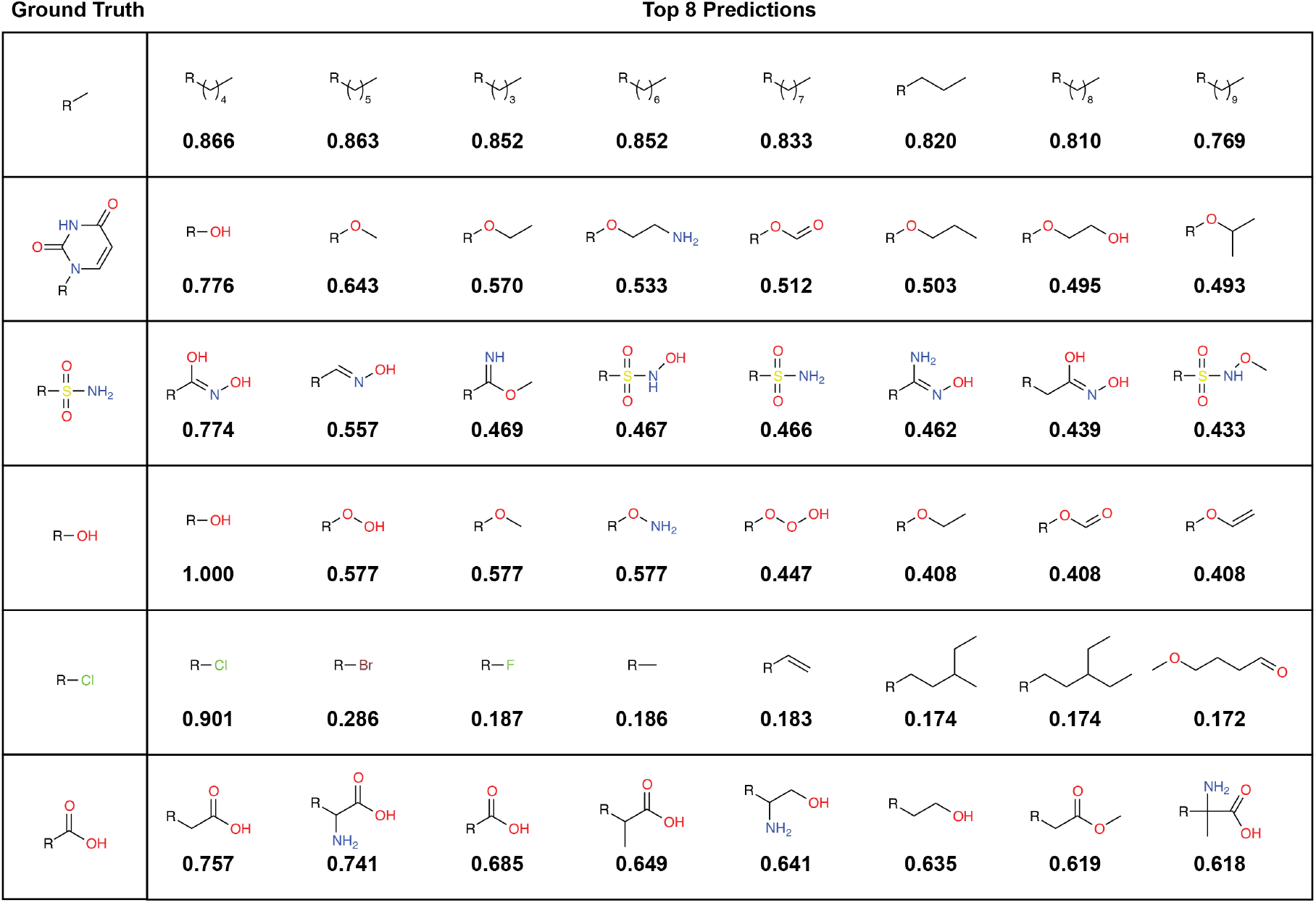
Example predictions using the final model. Examples are drawn from the (unseen) *TEST* set, and fragments are selected from the *LBL-ALL* label set (6,522 choices) Each predicted vector is the average of 32 predictions obtained by randomly rotating the corresponding input voxel grid. Left: The ground-truth (*correct*) fragments. Right: The top eight predicted fragments, labeled with the cosine similarity to the respective averaged prediction vector.

#### 4.5.2 Human and DeepFrag Intuition are Complementary

We also provide a simple comparison of human and DeepFrag “intuition” by considering the structure of *H. sapiens* peptidyl-prolyl cis-trans isomerase NIMA-interacting 1 (*Hs*Pin1p), a cancer drug target [26], bound to a phenyl-imidazole ligand (IC_50_ = 8 μM, PDB 2XP9 [27], Figure 5). Importantly, neither *Hs*Pin1p nor the ligand were included in the DeepFrag *TRAIN* or *VAL* sets.

**Figure 5:**
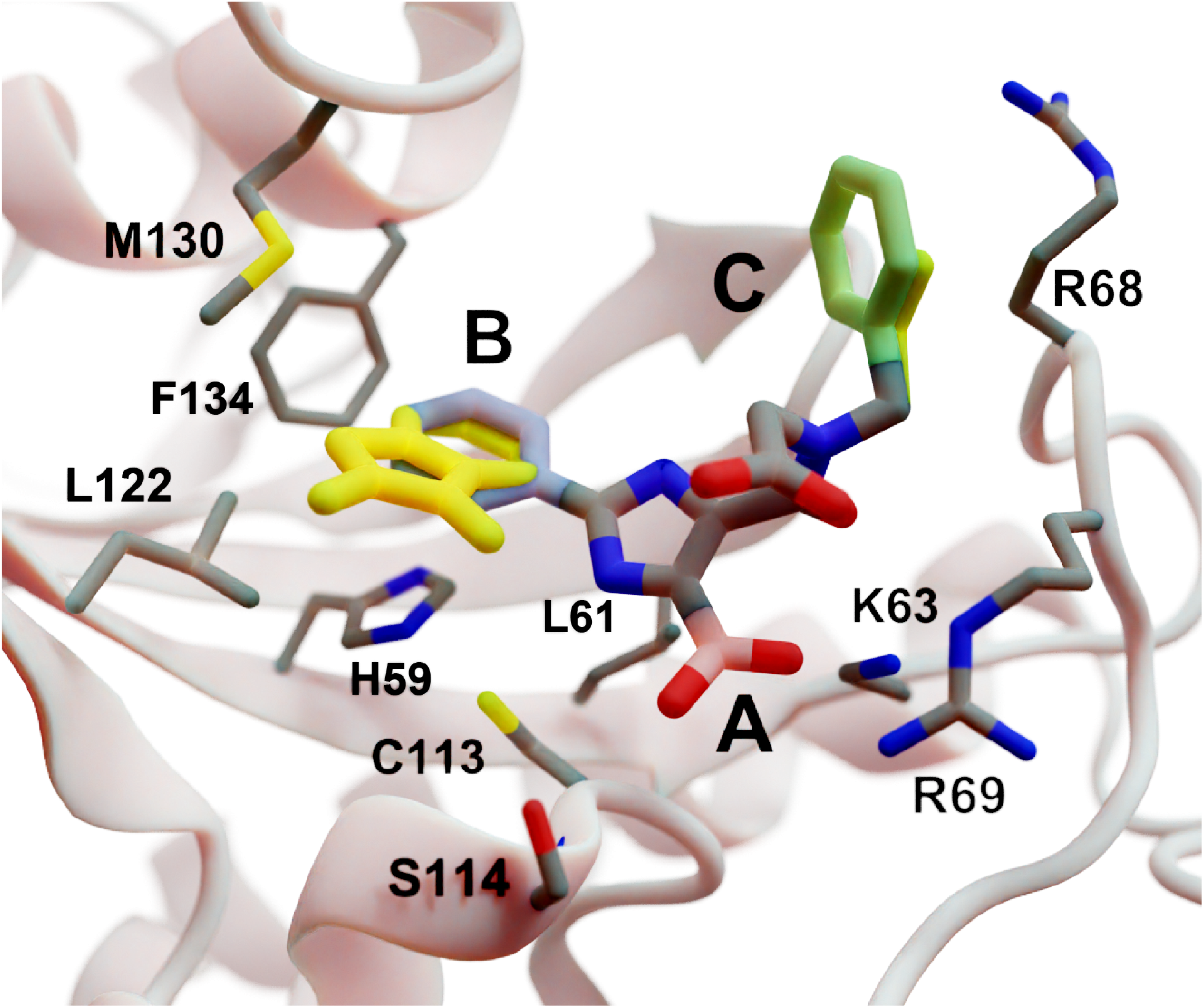
A crystal structure of *Hs*Pin1p bound to a phenyl-imidazole ligand (PDB 2XP9 [27]). A, B, and C indicate a carboxyl (pink) and two phenyl (blue and green) fragments that we reassessed with DeepFrag. The DeepFrag-suggested bicyclic and ethyl replacements for phenyl B and phenyl C, respectively, are shown in yellow.

Intuitively, carboxylate A (Figure 5, highlighted in pink) seems well optimized, per the crystal structure. It forms electrostatic interactions with R69 and K63, and hydrogen bonds with C113 and S114. We removed this carboxylate group and used DeepFrag to predict replacement moieties. The top predicted fragment was the correct carboxylate group. Interestingly, the second- and third-place fragments were chemically similar: *CO and *CC(=O)O.

Phenyl B (Figure 5, highlighted in blue) also appears to be well optimized. It binds near multiple hydrophobic residues (L122, F134, M130, and L61) and forms π-π interactions with H59. We repeated the same DeepFrag analysis multiple times, this time removing phenyl B. Multiple runs are useful in some cases because DeepFrag is not strictly deterministic; it randomly rotates the 32 grids it uses for output-fingerprint averaging, so different DeepFrag runs can in some cases predict different outputs. While some of our DeepFrag runs targeting phenyl B did identify a phenyl group as the *correct* fragment, we here describe a run in which the top predicted fragment was the bicyclic moiety *c1ccnc2c1C(=O)N(C)C2. To further examine the potential binding pose of this fragment, we used RDKit [21] to generate multiple fragment conformers/rotamers and to identify a parent-connected fragment position that was low-energy, per the UFF force field (Figure 5, in yellow) [28]. This fragment pose preserves the π-π interactions with H59 but potentially enhances hydrophobic interactions with M130 and L122, illustrating how DeepFrag can serve as a useful tool for lead optimization.

Finally, human intuition suggests that phenyl C (Figure 5, highlighted in green) is not well optimized. Aside from possible hydrophobic interactions with a portion of the R68 side chain, there are no other specific interactions with *Hs*Pin1p. This phenyl group also appears to be more solvent exposed than is phenyl B. When we applied DeepFrag, it in fact did not suggest aromatic groups at this position. The top predicted fragments were methyl and ethyl groups. To further examine the potential binding pose of the suggested ethyl moiety, we again identified a low-energy, parent-connected fragment pose (Figure 5, in yellow). Interestingly, this predicted pose maintains the potential hydrophobic interactions with the R68 side chain.

This analysis suggests that DeepFrag has learned much of the same chemical-biology intuition typical of experts in the field. The Supporting Information includes three additional examples of DeepFrag use.

### 4.6 Comparison with Other Approaches

Recently, machine-learning techniques have been applied to many tasks in computer-aided drug discovery. Specifically, the use of 3D-voxel descriptors in conjunction with convolutional neural networks is now commonplace, with reported applications for binding-affinity prediction [1–3], virtual screening [4–7], and QSAR [8].

Generative modeling appears to be a more challenging task. Several authors have repurposed generative architectures used in computer vision for use in drug design (e.g., generative adversarial networks, GANs; variational autoencoders, VAEs). For example, Skalic et al. developed a generative adversarial model derived from BicycleGan [29] that can create 3D pharmacophoric maps from a receptor target. A subsequent step generates SMILES strings using a recurrent “captioning network” [11]. Several other authors have created models that generate molecular analogues independently of a target receptor using a continuous, learned latent space [30, 31], generative recurrent networks [32], or deep reinforcement learning [33]. While promising, these approaches are difficult to evaluate quantitatively, making it challenging to decide when to use them in a production pipeline. In contrast, DeepFrag in unique in that it (1) formulates small-molecule lead optimization as a *classification* problem rather than a *generative modeling* problem and (2) predicts a fragment fingerprint given a 3D-voxel representation of a ligand/protein complex. To the best of our knowledge, ours is the first system to perform data-driven lead optimization in this way.

Others have applied more traditional approaches to fragment-based lead optimization that do not leverage machine learning. For example, a recent publication by Shan et al. [34] proposed *FragRep*, a program that also aims to guide local fragment optimization in the context of a binding pocket. Whereas *FragRep* uses hand-crafted rules to identify suitable fragments, *DeepFrag* infers these rules from a large dataset of examples. Additionally, generating a prediction in *DeepFrag* takes roughly 0.3 seconds on a GPU, compared to 60-120 seconds for *FragRep*. On the other hand, *FragRep* is advantageous in that it can also generate predictions for internal scaffold fragments (i.e. non-terminal fragments).

## 5 Conclusions

Lead optimization is a critical early step in the drug-discovery process. DeepFrag, a free machine-learning program aimed at guiding this important process, will thus be a useful tool for the computational-biology community. Though not a substitute for a trained medicinal chemist, DeepFrag is highly effective for hypothesis generation. It provides fragment suggestions that trained chemists and biologists can then evaluate, synthesize, and experimentally test.

Although DeepFrag accuracy is impressive, our approach has several notable limitations. Two of these limitations stem from our use of crystallographic data for training. First, crystallographic artifacts (e.g., due to crystallographic packing [35]) occasionally produce ligand/receptor conformations that differ from the physiological conformations that are most useful for drug discovery. Second, even when a crystallographic conformation is physiologically relevant, it represents only a single conformation. In reality, binding pockets are often highly dynamic, sampling multiple druggable conformations. And ligand/fragment binding can influence those dynamics via conformational-selection and induced-fit mechanisms [36]. Our approach also assumes that adding a fragment does not substantially impact the binding geometry of the initial parent portion of the ligand. This same assumption underlies many lead-optimization approaches, but it is possible that in some cases useful fragment additions might fundamentally alter the binding mode of the entire ligand. Our approach will likely fail in these cases.

Finally, to mitigate the challenges associated with ambiguous protonation states, the current version of DeepFrag simply ignores hydrogen atoms and makes predictions based on heavy-atom positions alone. We found that including hydrogen atoms leads to only modest improvements in accuracy (Table 4), but future computational methods that can accurately predict ionization and tautomerization may enable improved DeepFrag models.

These limitations aside, we expect DeepFrag to be useful in many applications. To encourage broad adoption, we release the DeepFrag model and software under the permissive Apache License, Version 2.0 (http://durrantlab.com/deepfragmodel). The git repository also includes a link to a Google Colaboratory Notebook [14] for testing.

## 6 Acknowledgements

We acknowledge Matthew Ragoza, Tomohide Masuda, and Dale Erikson for useful discussions. We also thank the University of Pittsburgh’s Center for Research Computing for providing helpful computer resources. This work was supported by the National Institute of General Medical Sciences of the National Institutes of Health [R01GM132353 to J.D.D. and R01GM108340 to D.R.K]. The content is solely the responsibility of the authors and does not necessarily represent the official views of the National Institutes of Health.

## Supporting Information

### 6.1 Hyperparameter Tuning

Our early efforts were focused on identifying model and grid-density parameters well suited to the fragment-selection task. Because this phase of hyperparameter tuning required us to generate many models for comparison, we considered only the receptor/ligand complexes present in the PDBBind database [37, 38], rather than those in the larger Binding MOAD database [15]. In this sense our early hyperparameter-tuning steps differed than those described in the main text. We note that the ligand SMILES strings in the PDBBind database are somewhat inconsistent (e.g., carboxylate moieties are inconsistently protonated), so that some fragment labels were duplicated in this dataset. This duplication led to overall lower accuracy. Fortunately, for the purpose of hyperparameter tuning we were concerned only with the relative performance between models. To further reduce the size of the dataset, we also considered only (*receptor/parent*, *fragment*) tuples with connection points that came within 3 Å of any receptor atom.

We divided these tuples into *TRAIN*, *TEST*, and *VAL* sets (60/20/20), using the same approach described in the main text. We trained on examples in the *TRAIN* set and evaluated TOP-k accuracy on examples in the *VAL* set. We merged the fragment vectors from the *TRAIN* and *VAL* sets into a combined label set that we used for fragment selection. We trained each model variant for 8 hours on a GPU, achieving accuracy roughly 80-90% of the eventual maximum.

In the first phase of early-stage hyperparameter tuning, we used a random search to identify reasonable model parameters. We randomly sampled 32 combinations of learning rates, batch sizes, grid widths, grid resolutions, and model architecture parameters (blocks and fc) (Table S1), where “blocks” describes the number of filters in each 3-part convolution block, and “fc” describes the number and sizes of fully-connected layers between the “Flatten” layer and the fragment output. For all models generated during this phase, we used an atom-influence radius of 2.0 Å, a grid resolution of 1.0 Å, the *SMOOTH* voxelation method, the *SUM* accumulation type, and the *simple* grid-layer definition for both the parent and receptor.

**Table S1:** Hyperparameters varied during our early-stage tuning protocol. Parentheses indicate optimal values that were fixed for later phases.

| Phase 1: Model parameters |  |
| --- | --- |
| Learning rate | 0.01, 0.001, **(0.0001)** |
| Batch size | **(16)**, 32, 64, 128 |
| Grid width | 16, **(24)**, 32 |
| Grid resolution | 0.5 Å, 1.0 Å, 2.0 Å |
| Blocks | [32,64], [16,32], [64,128], **([64,64])** |
| Fully Connected (fc) | [256], **([512])**, [1024], [2048], [256,256], [512,512], [1024,1024] |
| Phase 2: Grid-density parameters |  |
| Grid resolution | 0.25 Å to 3.00 Å **(0.75 Å)** |
| Atom-influence radius | 0.25 Å to 4.00 Å **(1.0-2.0 Å)** |

**Figure S1:**
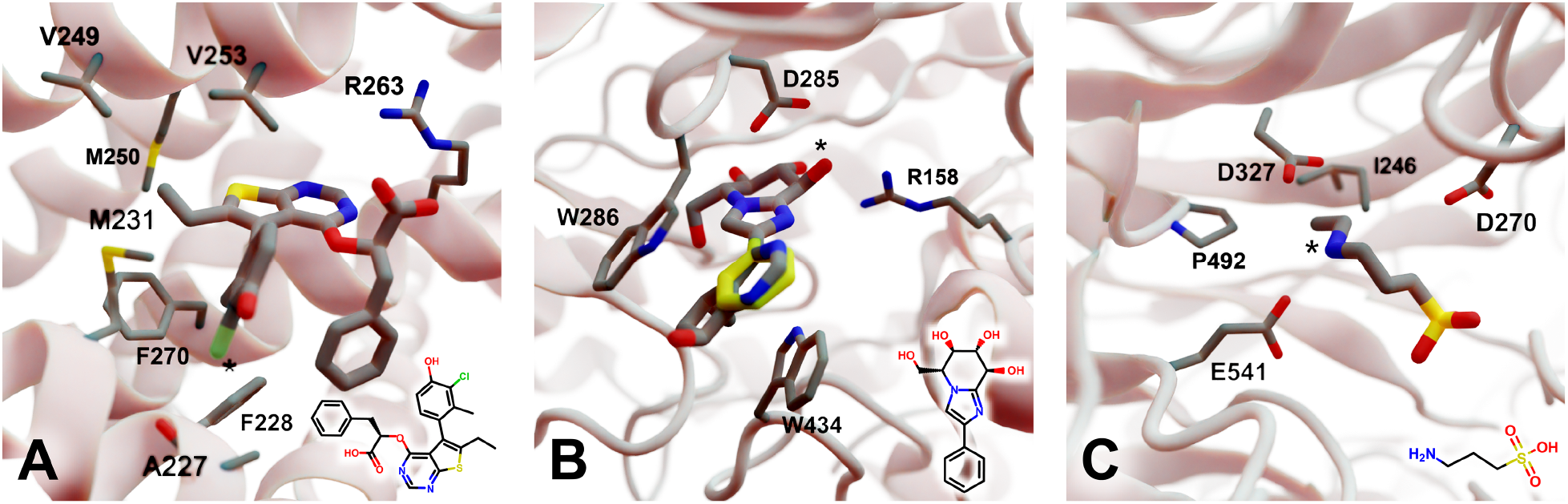
Crystal structures used to explore the DeepFrag model. The corresponding crystallographic ligands are overlaid in the lower right-hand corners. (A) Protein myeloid cell leukemia1 (Mcl-1) (PDB ID: 6QZ8 [39]). A key ligand chlorine atom is marked with an asterisk. (B) Family GH3 *β*-D-glucan glucohydrolase from barley (PDB ID: 1X38 [40]). A key ligand hydroxyl group is marked with an asterisk, and an original phenyl group is shown in yellow. (C) NanB Sialidase from *S. pneumoniae* (PDB ID: 4FOW [41]). A key ligand primary amine, which DeepFrag replaced with an ethylamine, is marked with an asterisk.

In the second phase of hyperparameter tuning, we fixed the best learning rate, batch size, grid width, and model architecture parameters found during the first phase. We then randomly sampled 32 different combinations of grid resolutions and atom-influence radii (Table S1). We continued to tune the grid resolution in the second phase because the first phase did not reveal an optimal value. All other parameters were the same as the first phase.

### 6.2 Additional Examples of Use

We here provide three additional examples of DeepFrag use. We selected these protein/ligand complexes because (1) the associated PDBs were among those in the *TEST* set; (2) the associated ligands had diverse low-weight fragments of the type commonly considered during lead optimization; and (3) visual inspection confirmed that those fragments formed specific interactions with their respective protein receptors, suggesting they are well optimized.

#### 6.2.1 Myeloid Cell Leukemia 1 (Mcl-1)

We first applied DeepFrag to the cancer-implicated protein myeloid cell leukemia 1 (Mcl-1) bound to a ligand designated “10d” (Figure S1A; PDB ID: 6QZ8 [39]). As a demonstration of DeepFrag’s ability to optimize for electrostatic interactions, we removed the ligand carboxylate group, which forms a strong electrostatic interaction with R263, and used DeepFrag to predict appropriate replacement fragments. The top predicted fragment was in fact a carboxylate group, and the second- and third-ranked fragments were chemically similar: *C(=O)OC and *C(=O)OO.

To demonstrate DeepFrag’s ability to optimize for hydrophobic interactions, we separately removed a terminal methyl and ethyl group from the ligand. Both appear to be well optimized. The methyl group forms hydrophobic contacts with F270, F228, and M231; and the ethyl group forms hydrophobic contacts with F270, V253, V249, M250, and M231. In both cases, DeepFrag identified the “correct” fragment as the top-ranked candidate and also suggested other chemically plausible hydrophobic fragments.

We also removed the ligand chlorine atom, which is predicted to form a halogen bond with the A227 backbone carbonyl oxygen atom (Figure S1A, marked with an asterisk). The top-ranked replacement fragments were methyl, methyl alcohol, and ethyl groups. Although methyl halide fragments did rank well (e.g., methyl floride, methyl chloride, methyl bromide, and methyl iodide ranked 8th, 12th, 13th, and 15th, respectively), it is reasonable that DeepFrag preferred a small, hydrophobic fragment (methyl) at this location because surrounding amino acids (i.e., F228, A227, and M231) are also hydrophobic.

#### 6.2.2 Family GH3 *β*-D-glucan Glucohydrolase (Barley)

We next applied DeepFrag to family GH3 *β*-D-glucan glucohydrolase from barley, bound to glucophenylimidazole (Figure S1B; PDB ID: 1X38 [40]). To show how DeepFrag can optimize for hydrogen-bond interactions, we first removed the hydroxyl group at position 8 (Figure S1B, marked with an asterisk), which participates in hydrogen bonds with R158 and D285. The top-ranked replacement fragment was the “correct” hydroxyl group, and other top-scoring fragments were chemically similar (e.g., *OC and *COO).

To test DeepFrag’s ability to optimize for aromatic stacking interactions, we next removed the ligand phenyl group (Figure S1B, in yellow), which participates in *π*-*π* stacking interactions with W286 and W434. Phenyl groups are larger than those tested above, making it less likely that DeepFrag will select the exact, “correct” fragment. But DeepFrag does often produce reasonable alternatives in these cases, showing that it has learned a certain degree of chemical intuition. The top replacement fragments were in fact all aromatic. The top fragment, *c1ncnc2c1C(C)CC2(O), was particularly interesting. To further examine its potential binding pose, we used RDKit [21] to generate multiple fragment conformers/rotamers and to identify a low-energy, parent-connected fragment position, per the UFF force field [28]. This pose suggests that the fragment’s bicyclic structure may enable more extensive contacts with W286 (Figure S1C). Interestingly, the fragment also overlaps with a crystallographic glycerol molecule (not shown), suggesting the expanded ligand may now occupy a new but “druggable” subpocket.

#### 6.2.3 NanB Sialidase (*Streptococcus pneumoniae*)

Finally, we applied DeepFrag to NanB Sialidase from *S. pneumoniae*, bound to 3-ammoniopropane-1-sulfonate (Figure S1C; PDB ID: 4FOW [41]). The primary amine of the ligand (marked with an asterisk) is likely positively charged because it is positioned among three negatively charged amino acids (E541, D327, and D270). To further illustrate how DeepFrag can account for electrostatic interactions, We removed the amino group and used DeepFrag to replace it. Though the “correct” amino group was among the top-ranked fragments (8th), the top fragment was in fact ethylamine (*NCC). We again used RDKit to find a low-energy, parent-connected fragment pose. The secondary amine of the expanded fragment could still form electrostatic interactions, but the expanded ethyl group may form additional hydrophobic interactions with I246 and P492, possibly explaining why DeepFrag prefered the larger fragment at this position.

